# Socioeconomic Differentials in Hypertension based on JNC7 and ACC/AHA 2017 Guidelines Mediated by Body Mass Index: Evidence from Nepal Demographic and Health Survey

**DOI:** 10.1101/667899

**Authors:** Juwel Rana, Zobayer Ahmmad, Kanchan Kumar Sen, Sanjeev Bista, Rakibul M Islam

## Abstract

**Background:** Unlike developed countries; higher socioeconomic status (SES, education, and wealth) is associated with hypertension in low and middle-income countries (LMICs) with limited evidence. We examined the associations between SES and hypertension in Nepal and the extent to which these associations vary by sex and urbanity. The body mass index (BMI) was examined as a secondary outcome and assessed as a potential mediator.

**Materials and methods:** We analyzed the latest Nepal Demographic and Health Survey data (N=13,436) collected between June 2016 and January 2017, using a multistage stratified sampling technique. Participants aged 15 years or older from selected households were interviewed with an overall response rate of 97%. Main outcomes were hypertension and normal blood pressure defined by the widely used Seventh Report of the Joint National Committee (JNC 7), and the American College of Cardiology/American Heart Association (ACC/AHA) 2017.

**Results:** The prevalence of hypertension was higher in Nepalese men than women. The likelihood of having hypertension was more than double for individuals in the highest versus lowest wealth quintiles [men: OR 2.13, 95% CI 1.60-2.85); women: OR 2.54, 95% CI 2.00- 3.24] and for individuals with the higher education versus no education [men: OR 2.38, 95% CI 1.75-3.23; women: OR 1.63, 95% CI 1.18-2.25]. The associations between SES and hypertension were different by sex and urbanity. These associations were mediated by BMI.

**Conclusions:** Higher SES was positively associated with the higher likelihood of having hypertension in Nepal according to both JNC 7 and ACC/AHA 2017 guidelines. These associations were mediated by BMI, which may help to explain broader socioeconomic differentials in CVD and related risk factors, particularly in terms of education and wealth. Our study suggests that the mediating factor of BMI should be tackled to diminish the risk of CVD in people with higher SES in LMICs.

## Introduction

Hypertension is a growing public health problem in low and middle-income countries (LMICs)^1^ with concurrent risks of cardiovascular and kidney diseases.^2^ A review warned that although about three-quarters of people with hypertension (639 million people) live in LMICs, there is no improvement in awareness or control rates.^1^ Hypertension is a major contributor to death and disability in South Asian countries, including Nepal with a low level of control and awareness.^3–6^ The World Health Organization (WHO) implemented `STEP-wise approach to surveillance’ (STEPS) using nationally representative sample in 2008 and 2013 reported an increasing trend of prevalence of hypertension among 15-69 years Nepalese population ranging from 21.5% in 2008 to 26.0% in 2013.^7,8^ Based on the recent Nepal Demographic and Health Survey (NDHS) 2016, Kibria and colleagues reported that the estimated prevalence of hypertension in Nepal using the widely used Seventh Report of the Joint National Committee (JNC 7) guideline^9^ was 21.2%, and the corresponding prevalence was 44.2% when using a new hypertension guideline recommended by the American College of Cardiology/American Heart Association 2017 (ACC/AHA 2017).^10^ This study demonstrated that the prevalence of hypertension increased to 23% when using new ACC/AHA guideline, with the highest increase in the richest and obese population.^11^

Despite an increasing prevalence of hypertension in Nepal, research exploring complex interrelationship between socioeconomic status (SES), indicated by education levels and wealth quintiles, and hypertension is limited. Moreover, this association is complex, unlike developed countries, in LIMCs. For instance, the prevalence of hypertension is higher among low SES group in developed countries,^12,13^ while it is substantially higher among high SES groups in LMICs.^14,15^

The reasons for the high prevalence of hypertension in the low SES group in developed countries include higher smoking rates, higher body mass index (BMI), and lack of exercise compared with higher SES groups.^16^ The opposite pattern is observed in LMICs, where a higher prevalence of these risk factors is observed in higher SES groups compared with low SES group. A recent review found that people in higher SES groups in LMICs were less likely to be physically active and consume more fats, salt, and processed food than low SES group.^17^ Furthermore, studies also found that BMI is exponentially increasing in people in LIMCs,^18–20^, which are the key modifiable risk factors for hypertension. Thus, we hypothesized that there would be positive associations between high SES and hypertension in Nepal, and the level of BMI will at least partially mediate these associations. The primary aim of this study was (i) to assess the associations between SES and hypertension in Nepal, and the extent to which these associations vary by gender and urbanity; and (ii) to examine whether BMI attenuates the associations between SES and hypertension and, the extent to which BMI explain these associations. The secondary aim of this study was to examine associations of BMI with SES and the extent to which these associations vary by gender and urbanity.

## Materials and Methods

### Data source

The study analyzed the nationally representative Nepal Demographic and Health Survey (NDHS) 2016 data, collected between June 2016 and January 2017. The Nepal Health Research Council and the ICF International institutional review board approved the NDHS 2016 survey protocol. The household head provided written informed consent before the interview. For the current study, we obtained approval to use the data from ICF in June 2018.

### Survey design and study populations

The updated version of the census frame of National Population and Housing Census 2011 conducted by the Central Bureau of Statistics was used as the sampling frame for the NDHS 2016. The households of the NDHS 2016 were selected in two ways based on the urban/rural locations. Firstly, the two-stage stratified sampling process was used in rural areas where wards were selected in the first stage as a primary sampling unit (PSUs) and households were selected in the second stage. Secondly, three-stage stratified sampling was used in urban areas to select households where wards were selected in the first stage (PSUs), enumeration areas (EA) were selected from each PSU in the second stage and households were selected from EAs in the third stage. There were 14 sampling strata in the NDHS 2016, where wards were selected randomly from each stratum. A total of 383 wards were selected altogether, 184 from urban and 199 from rural areas. Finally, a total of 11,490 households (rural- 5,970 and urban-5,520) were selected for the NDHS 2016.^21^ Flowchart of the analytic sample selection process is given as supplementary figure (S1 Fig.).

The trained interviewers collected data visiting the households. The overall response rate was approximately 97%. Blood pressure (BP) was measured among 15,163 individuals with 6,394 men and 8,769 women aged 15 years and above. In our study, a total unweighted sample was 13,371 comprising men (5,535) and women (7,836), after excluding participants aged <18 years and discarding the missing and extreme values. The total weighted analytic sample was 13,436 participants (men 5,646 and women 7,790) aged 18 years and above. Details of the NDHS 2016, including survey design, sample size determination, and questionnaires have been described elsewhere.^21^

### Measures of outcomes: Blood Pressure Outcomes

Hypertension and normal blood pressure were considered as the outcome variables in the study defined by both JNC 7^9^ and ACC/AHA 2017 guidelines (Table 1).^10^ Three measurements of blood pressure (systolic and diastolic blood pressures) were taken for each participant with an interval of 5 minutes between the measurements by UA-767F/FAC (A&D Medical) blood pressure monitor. Systolic blood pressure (SBP) and diastolic blood pressure (DBP) were defined by taking the average of three SBP and three DBP measurements, respectively. We used both ‘measurement-only’ and ‘medical/clinical’ definitions to generate independent binary outcomes for ‘hypertension’ and ‘normal blood pressure’ based on both guidelines. The ‘measurement-only’ definition was developed solely based on the cut-off points that accounted for the average of three SBP and three DBP measurements. The ‘medical/clinical’ definition accounted for ‘measurement-only’ definition plus medical diagnosis by a health professional as having high blood pressure and or taking medication for lowering high blood pressure (Table 1).

**Table 1.** Definitions of Blood Pressure Outcome Used in the Study

| Blood pressure outcomes | Measurement-only definitions | Medical definitions |
| --- | --- | --- |
| Hypertension (JNC 7)<sup>†</sup> | SBP ≥140mmHg or DBP ≥ 90 mmHg | Meet any of the following three criteria:<br>(1) SBP ≥ 140mmHg or DBP ≥ 90mmHg<br>(2) Doctor/nurse diagnosed high blood pressure<br>(3) Taking blood pressure-lowering medication |
| Hypertension (ACC/AHA 2017)<sup>‡</sup> | SBP ≥130mmHg or DBP ≥ 80 mmHg | Meet any of the following three criteria:<br>(1) SBP ≥ 130mmHg or DBP ≥ 80mmHg<br>(2) Doctor/nurse diagnosed high blood pressure<br>(3) Taking blood pressure-lowering medication |
| Normal blood pressure (JNC 7 or ACC/AHA 2017) | SBP <120mmHg and DBP <80 mmHg | SBP ≤ 120mmHg and DBP ≤ 80 mmHg, no diagnosis of high blood pressure, and not taking blood pressure-lowering medication |

†JNC 7= The Seventh Report of the Joint National Committee
‡ACC/AHA 2017= The 2017 American College of Cardiology/American Heart Association

### Measures of Exposure: Socioeconomic status

Three indicators such as education levels, wealth quintiles, and employment status are most commonly used in several studies to assess the SES of a participant,.^12, 15^ However, we omitted employment status from our assessment of SES and subsequent analyses due to a large number of missing values as the majority of the women in South Asia are not involved in formal employment. The NDHS 2016 provided data for a derived wealth quintile using the principal component analysis taking scores of a household’s durable and nondurable assets. Firstly, households are given scores using principal component analysis based on the number and kinds of consumer goods they own. Secondly, to get the wealth quintiles, the distribution of scores was divided into five equal categories named as poorest, poorer, middle, richer, and richest. Education was an ordinal measure of self-reported levels of education, which was grouped into four different categories (no education/preschool, primary, secondary, and higher education) in the NDHS 2016. In our study, the SES measures were not indexed for two main reasons. Firstly, different indicators of SES tend to have different theoretical pathways to BMI and blood pressure outcomes. Secondly, SES indicators might be causally related to each other; and they build on each other according to the life course models.^22^

### Body Mass Index

The BMI was used in the study as both continuous and categorical variables. We followed both the South-Asian specific and global definition of BMI.

### Statistical analysis

Our primary statistical analyses assessed the sex and urbanity stratified associations of educational levels and wealth quintiles with blood pressure outcomes using both the measurement-only and medical definitions (Table 1; Fig 1–2; S1-4 Tables; S2-4 Fig.). To characterize the shapes of the associations, we calculated sex and urbanity stratified adjusted odds ratios (ORs) and 95% confidence intervals (CIs) within each level of education or wealth quintiles by using the binary logistic regression models. We used a cutoff of 10% change in the stratified analysis to identify differences in hypertension by sex and urbanity.

**Fig 1:**
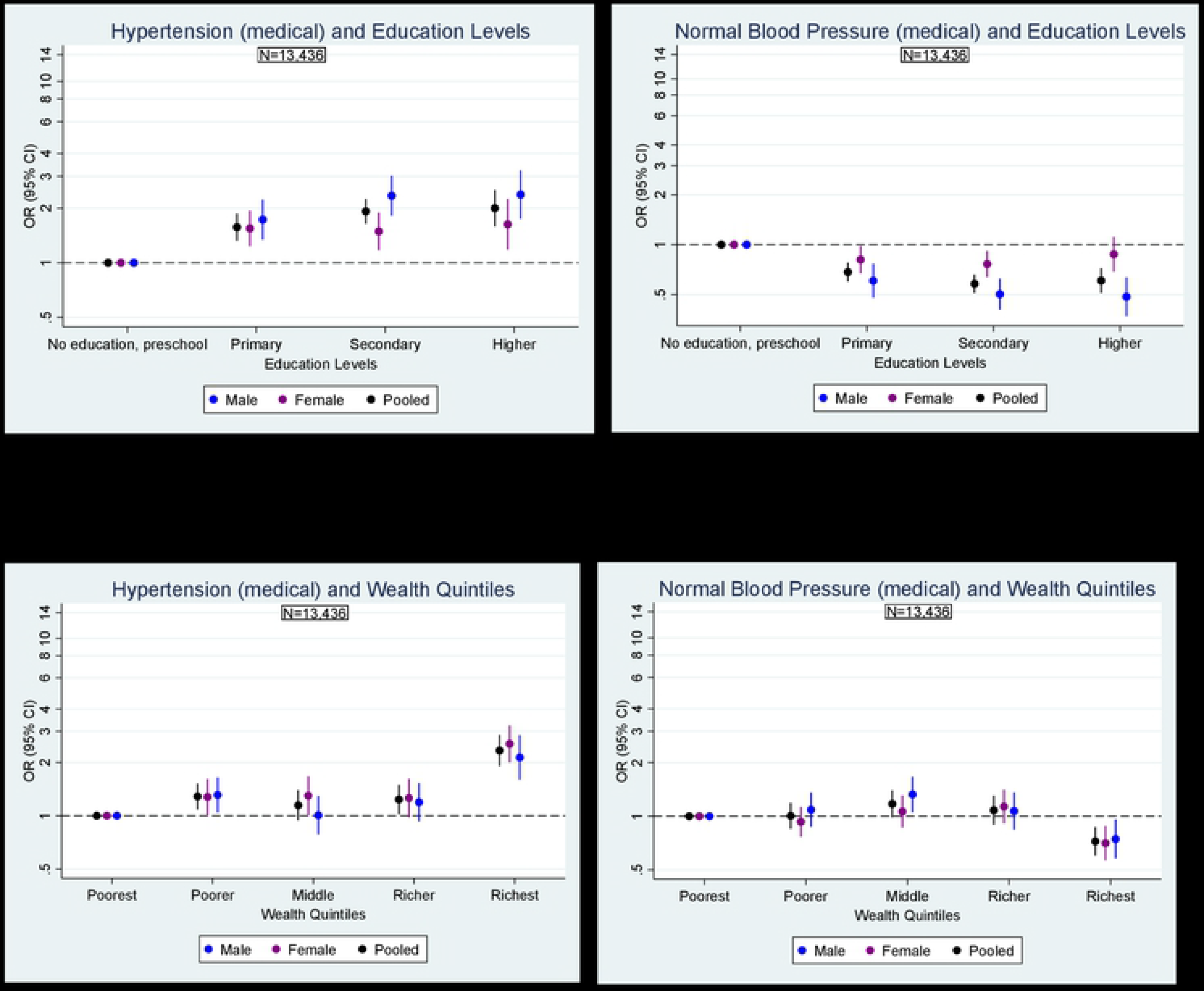
Association of blood pressure outcome (medical) with education levels by sex in Nepal (In a table format as supplementary data, S1 Table) Odds ratios are adjusted for age, urbanity and marital status, and stratified by sex. Medical outcomes are defined based on cut-off points, diagnosis by a health professional, or relevant medication use. Cut-off points are defined as follows: hypertension: SBP ≥140mmHg or DBP ≥90mmHg; normal blood pressure: SBP ≤ 120mmHg and DBP ≤ 80 mmHg. Measurement-only outcomes are defined based on cut-off points only and the association of blood pressure outcome (measured) with education levels by sex in Nepal.

We further tested whether BMI mediates the associations between SES and blood pressure outcomes (medical). There were two general approaches to testing mediation models. The first approach was the “reduction-in-estimate criterion” approach (Table 4; S7 Table), which assessed whether the inclusion of mediator variable-BMI attenuated the associations or effects for the main predictors across nested models. Hence, we constructed two nested models, and coefficients were progressively adjusted for age, sex, marital status, urbanity, and second-hand smoking in model 1. Coefficients were further adjusted for prior determined mediator-BMI in model 2 to observe the changes in the effects of predictors. We considered that there is a mediation effect of BMI using a cutoff of 10% reduction in the effect estimate after adjusting for BMI in the model 2. The second step was the “indirect effect” approach, which formally examined the statistical significance of an indirect effect using the product of coefficients approach. For assessing the indirect effect of BMI on these binary outcomes, we used the binary-mediation package in Stata because this approach is commonly used and can detect which variables are continuous and which are binary. It requires information for each link in the proposed mediation process (X-M-Y). In supplemental analyses, we replicated the process for 15 000 bootstraps for statistical significance, which provided substantially identical indirect and direct effects along with standard errors and biased-corrected 95% CIs.

Additionally, we examined the adjusted associations between SES and BMI as a continuous outcome. Moreover, the adjusted sex and urbanity stratified associations between SES and binary outcome-overweight/obese (using both global and South Asia-specific cut-offs for BMI) were also assessed to observe differences in overweight/obesity by sex and urbanity.

To examine the associations between SES and hypertension, all potential confounders for each predictor were selected using prior knowledge and directed acyclic graphs (DAGs) to avoid the ‘Table 2 fallacy’ in a multivariable model and to observe unbiased total effect estimates for predictors.^23, 24^

**Table 2:** Sample characteristics (weighted numbers and percentages unless stated otherwise)

| Characteristics | Overall<br>(n=13 436) | Men<br>(n = 5645) | Women<br>(n= 7790) | p-value |
| --- | --- | --- | --- | --- |
| Mean Age (SE, standard error) | 40.7 (0.1) | 42.59 (0.28) | 39.30 (0.19) | < 0.001 |
| Marital Status (%) |  |  |  |  |
| Unmarried | 1569 (11.7) | 872 (15.4) | 698 (9.0) | < 0.001 |
| Married | 11 867 (88.3) | 4 774 (84.6) | 7092 (91.0) |  |
| Education Levels (%) |  |  |  |  |
| No education/preschool | 5498 (40.9) | 1474 (26.1) | 4024 (51.7) | < 0.001 |
| Primary | 2281 (17.0) | 1194 (21.2) | 1087 (14.0) |  |
| Secondary | 3709 (27.6) | 1958 (34.7) | 1751 (22.5) |  |
| Higher | 1947 (14.5) | 1020 (18.1) | 928 (11.9) |  |
| Employment Status (%) |  |  |  |  |
| Unemployed | 2777 (30.6) | 557 (15.9) | 2,220 (40.0) | < 0.001 |
| Employed | 6287 (69.4) | 2956 (84.15) | 3331 (60.0) |  |
| Wealth Index (%) |  |  |  |  |
| Poorest | 2405 (17.9) | 993 (17.6) | 1412 (18.1) | < 0.001 |
| Poorer | 2613 (19.5) | 1054 (18.7) | 1559 (20.0) |  |
| Middle | 2693 (20.0) | 1091 (19.3) | 1603 (20.6) |  |

For the brevity, we have reported ‘medical definition’ of hypertension/normal blood pressure, if not stated otherwise, especially when we assessed associations between hypertension/normal blood pressure and SES. However, similar analyses for ‘measurement only’ outcome of hypertension have been provided as supplementary data. Comparable analyses based on the new guideline of ACC/AHA 2017 have also been given as supplementary data. Two-sided *P*- values and 95% CIs are presented. The complex survey design effects were accounted in all performed analyses for reducing differences due to oversampling, variation in the probability of selection and non-response in the NDHS 2016. All analyses were performed using Stata 15 (StataCorp).

## Results

### General characteristics of study participants

Of 13,436 participants, 7,790 (58%) were women, and 5,645 (42%) were men, with a mean age of 40.7 (SE ±0.10) years (Table 2). More than half (61.1%) of the population lived in urban areas with no significant sex difference. About 40% of the population had no education, and men were more likely to be educated than women at each level of education (p <0.001). Men were also more likely to be wealthier than women were (p <0.0001).

|  |  |  |  |  |
| --- | --- | --- | --- | --- |
| Richer | 2936 (21.9) | 1280 (22.8) | 1656 (21.3) |  |
| Richest | 2787 (20.8) | 1228 (21.8) | 1559 (20.0) |  |
| Urbanity (%) |  |  |  |  |
| Urban | 8205 (61.1) | 3475 (61.6) | 4729 (60.7) | 0.27 |
| Rural | 5231 (38.9) | 2171 (38.6) | 3061 (39.3) |  |
| Region (%) |  |  |  |  |
| Mountain | 859 (6.4) | 367 (6.5) | 491 (6.3) | 0.60 |
| Hill | 5922 (44.1) | 2468 (43.7) | 3454 (44.3) |  |
| Teraï | 6655 (49.5) | 2811 (49.8) | 3844 (49.4) |  |
| Established Risk Factors of Hypertension |  |  |  |  |
| Mean Systolic Blood Pressure (SE) | 117.7 (0.2) | 122.02 (0.43) | 114.57 (0.38) | < 0.001 |
| Mean Diastolic Blood Pressure (SE) | 78.3 (0.1) | 79.89 (0.32) | 77.17 (0.26) | < 0.001 |
| High Blood Pressure (Told by doctor, %) | 1670 (12.4) | 763 (13.52) | 907 (11.64) | 0.004 |
| Medication for Blood Pressure (%) | 578 (4.3) | 260 (4.61) | 318 (4.08) | 0.25 |
| Mean Body Mass Index (SE) | 22.1 (0.0) | 21.08 (0.72) | 22.28 (0.10) | < 0.001 |
| Exposure to Secondhand Smoking (%) | 6308 (47.0) | 2718 (48.2) | 3589 (46.08) | 0.003 |
| Consumption of Caffeine (%) | 1058 (8.0) | 581 (10.3) | 477 (6.12) | < 0.001 |

A similar trend was found for employment status where men were about 24% higher in employment status than women were (p <0.001). Mean BMI was significantly higher among women (22.28 vs. 21.08; *p*< 0.001) compared with men. Men were more likely to be exposed to secondhand smoking (p <0.003) compared with women.

### Prevalence of hypertension by sex and urbanity

Women were having lower prevalence of hypertension compared with men for both measured (16.0%, 95% CI: 14.8, 17.3 vs. 22.8%, 95% CI: 21.2, 24.5) and medical hypertension (21.7%, 95% CI: 20.4, 23.0 vs. 29.1%, 95% CI: 27.4, 30.8) and the differences were significant statistically in both measurements (*p*< 0.001) (Table 3). People living in urban areas were having higher prevalence of hypertension compared with people living in rural areas for both measured (19.5%, 95% CI: 18.7, 20.4 vs. 17.9%; 95% CI: 16.9, 19.0) and medical (26.2%, 95% CI: 25.2, 27.1 vs. 22.7%; 95% CI: 21.6, 23.8) hypertension and the differences were significant statistically (*p*< 0.001) only for medical hypertension. Comparable trends were observed for both measurements in normal blood pressure (p <0.001). According to the new ACC/AHA 2017 guideline, there was an overall 21% increase of prevalence of hypertension, with the highest increase in the male population (23%). Similar trends of sex differences were observed in both hypertensions (p <0.001), however urban-rural significant differences (p >0.05) were not observed.

**Table 3:**
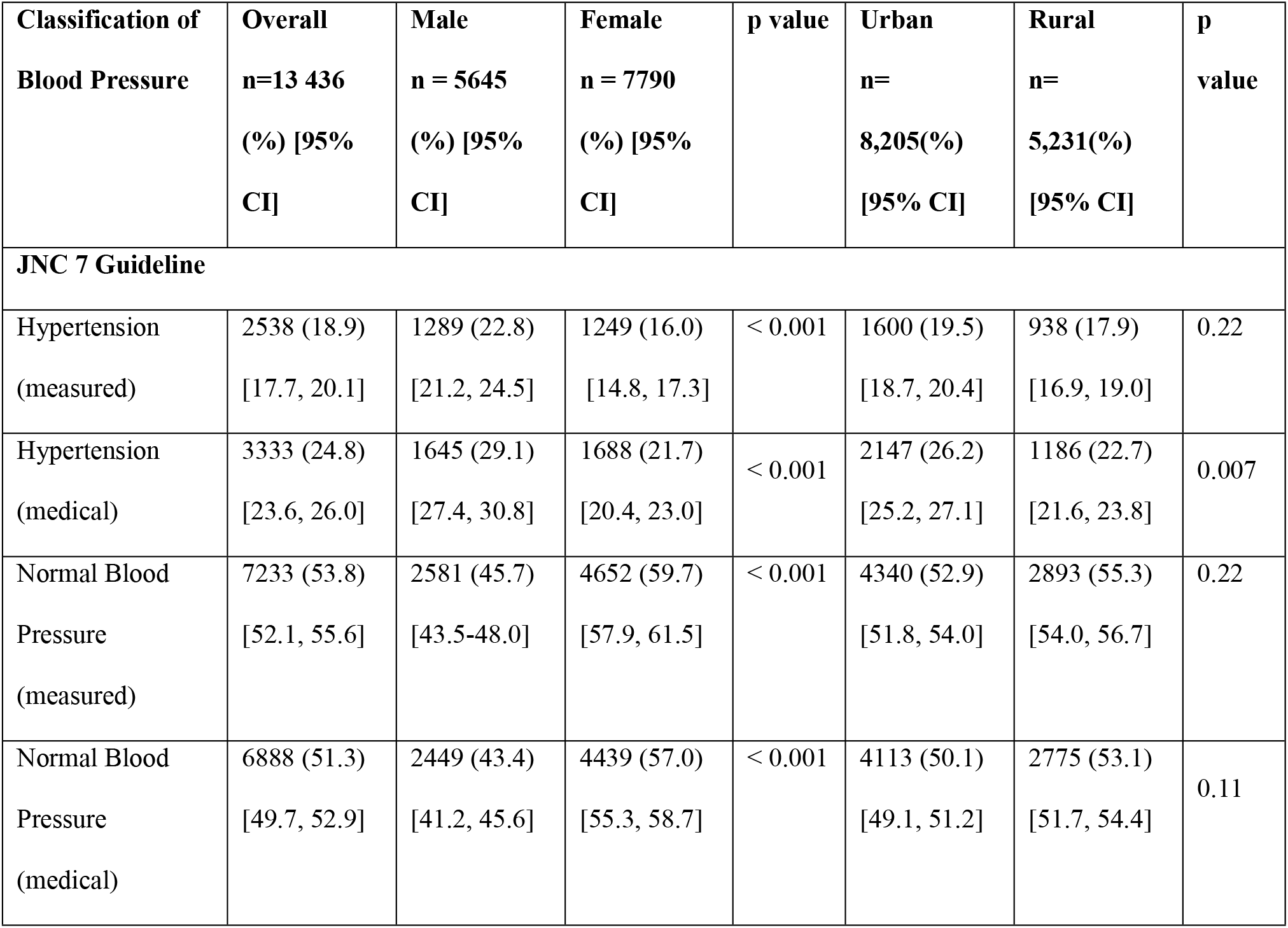
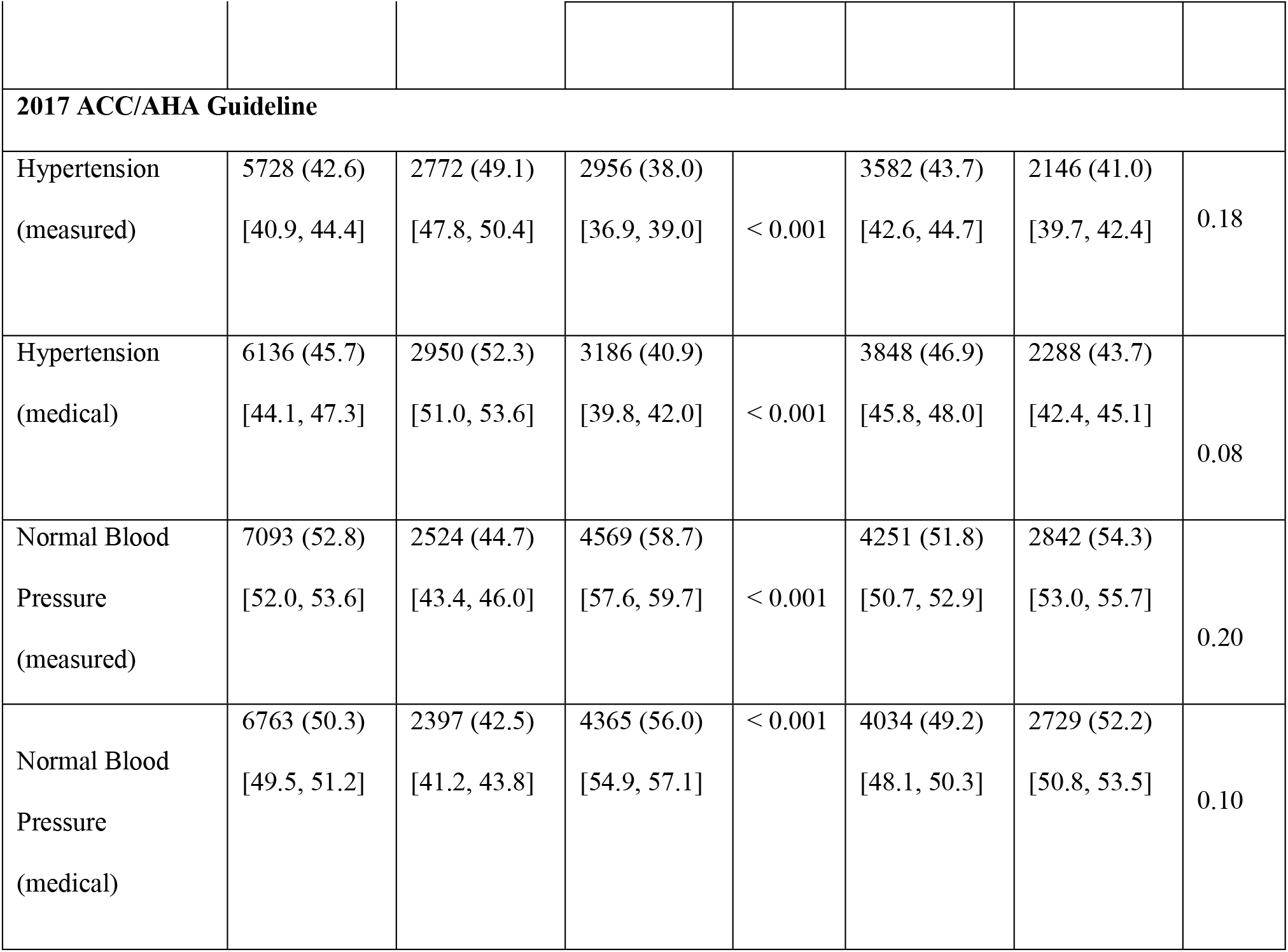
Prevalence of hypertension by sex and urbanity in Nepal

| Classification of Blood Pressure | Overall<br>n=13 436<br>(%) [95% CI] | Male<br>n = 5645<br>(%) [95% CI] | Female<br>n = 7790<br>(%) [95% CI] | p value | Urban<br>n=<br>8,205(%)<br>[95% CI] | Rural<br>n=<br>5,231(%)<br>[95% CI] | p value |
| --- | --- | --- | --- | --- | --- | --- | --- |
| **JNC 7 Guideline** |  |  |  |  |  |  |  |
| Hypertension (measured) | 2538 (18.9)<br>[17.7, 20.1] | 1289 (22.8)<br>[21.2, 24.5] | 1249 (16.0)<br>[14.8, 17.3] | < 0.001 | 1600 (19.5)<br>[18.7, 20.4] | 938 (17.9)<br>[16.9, 19.0] | 0.22 |
| Hypertension (medical) | 3333 (24.8)<br>[23.6, 26.0] | 1645 (29.1)<br>[27.4, 30.8] | 1688 (21.7)<br>[20.4, 23.0] | < 0.001 | 2147 (26.2)<br>[25.2, 27.1] | 1186 (22.7)<br>[21.6, 23.8] | 0.007 |
| Normal Blood Pressure (measured) | 7233 (53.8)<br>[52.1, 55.6] | 2581 (45.7)<br>[43.5-48.0] | 4652 (59.7)<br>[57.9, 61.5] | < 0.001 | 4340 (52.9)<br>[51.8, 54.0] | 2893 (55.3)<br>[54.0, 56.7] | 0.22 |
| Normal Blood Pressure (medical) | 6888 (51.3)<br>[49.7, 52.9] | 2449 (43.4)<br>[41.2, 45.6] | 4439 (57.0)<br>[55.3, 58.7] | < 0.001 | 4113 (50.1)<br>[49.1, 51.2] | 2775 (53.1)<br>[51.7, 54.4] | 0.11 |
| **2017 ACC/AHA Guideline** |  |  |  |  |  |  |  |
| Hypertension<br>(measured) | 5728 (42.6)<br>[40.9, 44.4] | 2772 (49.1)<br>[47.8, 50.4] | 2956 (38.0)<br>[36.9, 39.0] | < 0.001 | 3582 (43.7)<br>[42.6, 44.7] | 2146 (41.0)<br>[39.7, 42.4] | 0.18 |
| Hypertension<br>(medical) | 6136 (45.7)<br>[44.1, 47.3] | 2950 (52.3)<br>[51.0, 53.6] | 3186 (40.9)<br>[39.8, 42.0] | < 0.001 | 3848 (46.9)<br>[45.8, 48.0] | 2288 (43.7)<br>[42.4, 45.1] | 0.08 |
| Normal Blood<br>Pressure<br>(measured) | 7093 (52.8)<br>[52.0, 53.6] | 2524 (44.7)<br>[43.4, 46.0] | 4569 (58.7)<br>[57.6, 59.7] | < 0.001 | 4251 (51.8)<br>[50.7, 52.9] | 2842 (54.3)<br>[53.0, 55.7] | 0.20 |
| Normal Blood<br>Pressure<br>(medical) | 6763 (50.3)<br>[49.5, 51.2] | 2397 (42.5)<br>[41.2, 43.8] | 4365 (56.0)<br>[54.9, 57.1] | < 0.001 | 4034 (49.2)<br>[48.1, 50.3] | 2729 (52.2)<br>[50.8, 53.5] | 0.10 |

### Socioeconomic status and hypertension by sex and urbanity

Fig 1 and 2 explained the odds of blood pressure outcomes by education and wealth quintiles. The likelihood of being hypertensive (medical) was significantly higher in higher education group compared with lowest or no education group for men (OR 2.38 95% CI: 1.75, 3.23) and for women (OR 1.63 95% CI: 1.18, 2.25). People in the richest group were more likely to be hypertensive (medical) compared with people in the poorest group for men (OR 2.13 95% CI: 1.60, 2.85) and for women (OR 2.54 95% CI: 2.00, 3.24). Furthermore, the associations between SES and hypertension were significant and were different by sex and urbanity (S1-4 Tables). Similar trends and associations between hypertension and SES were observed by ACC/AHA 2017 guidelines (S2-4 Fig). Similarly, people with higher SES were less likely to have normal blood pressure compared with people in low SES (S1-4 Tables).

**Fig 2:** Association of blood pressure outcome (medical) with wealth quintiles by sex in Nepal (In a table format as supplementary data, S2 Table). Odds ratios are adjusted for age, urbanity and marital status, and stratified by sex. Medical outcomes are defined based on cut-off points, diagnosis by a health professional, or relevant medication use. Cut-off points are defined as follows: hypertension: SBP ≥140mmHg or DBP ≥90mmHg; normal blood pressure: SBP ≤ 120mmHg and DBP ≤ 80 mmHg. Measurement-only outcomes are defined based on cut-off points only and the association of blood pressure outcome (measured) with wealth quintiles by sex in Nepal.

### Mediation effect of BMI on SES and hypertension

Tables 4 shows a reduction in estimates of SES after including BMI in the logistic regression model. For the level of education, the adjusted odds of hypertension (medical) significantly decreased throughout the models, and particularly BMI attenuated the association and level of significance for each primary, secondary and higher education category with hypertension (medical) in the model 2 (Table 4: Model 1 vs. Model 2).

**Table 4:** Mediation effect of BMI on hypertension (medical) by education levels and wealth quintiles in Nepal

| Predictor | Model 1<sup>§</sup> | Model 2 |
| --- | --- | --- |
|  | Adjusted Beta coefficients<br>(95 % CI) | Adjusted Beta coefficients<br>(95 % CI) |
| **Education (Ref. No education/preschool)** |  |  |
| Primary | 0.40 (0.22, 0.58)*** | 0.24 (0.06, 0.42)* |
| Secondary | 0.58 (0.42, 0.75)*** | 0.35 (0.17, 0.53)*** |
| Higher | 0.61 (0.38, 0.85)*** | 0.32 (0.06, 0.57)* |
| **Wealth Quintiles (Ref. Poorest)** |  |  |
| Poorer | 0.25 (0.08, 0.42)** | 0.21 (0.04, 0.38)* |
| Middle | 0.13 (-0.07, 0.33) | 0.08 (-0.11, 0.27) |
| Richer | 0.20 (0.01, 0.39)* | 0.02 (-0.18, 0.21) |
| Richest | 0.84 (0.62, 1.05)*** | 0.39 (0.16, 0.62)*** |

<sup>§</sup>Coefficients adjusted for age, sex, marital status, urbanity, and second-hand smoking;
<sup>||</sup>Coefficients further adjusted for body mass index (BMI); Exponentiated coefficients; 95% confidence intervals in brackets
\* p<0.05, \*\* p<0.01, \*\*\* p<0.001.

Table 4 also suggests that further adjustment for BMI in model 2 reduced the effect size and level of significance in wealth quintiles and hypertension (medical). In other words, the inclusion of BMI in the model 2 has reduced the beta coefficient of hypertension for wealth quintiles and reduced statistical significance (Table 4: Model 1 vs. Model 2). BMI, therefore, may have a mediation effect on the association between SES and blood pressure outcomes. Similar analyses were also performed for normal blood pressure (medical) [S7 Table].

### Mediation Analysis: Body Mass Index

We formally tested the mediation effect of BMI on hypertension (medical) and SES as well as presented the unstandardized path coefficients (standard errors) and indirect effect (mediation) of BMI with bias-corrected 95% CI (Fig 3). A statistically significant mediation effect was observed for education (Coef. 0.04 95% CI: 0.03 to 0.05), which was substantially larger than the direct effect (Coef. −0.16 95% CI: −0.19 to −0.14). The proportion of the total effect that is mediated was about −0.31, which was also substantial. BMI also had mediation effect (Coef. 0.07 95% CI: 0.06 to 0.08) in the association between wealth quintiles and hypertension (medical), with 91% of the total effect (of wealth quintiles on hypertension) being mediated (by BMI). It was substantially higher than the direct effect (Coef. 0. 01 95% CI: −0.02 to 0.03).

**Fig 3:** Mediating Role of BMI in the Associations between SES and Hypertension (Medical) in Nepal Unstandardized path coefficients (standard errors) and unstandardized indirect effect (mediation) of BMI with bias-corrected 95% confidence intervals are reported.

### Sensitivity analyses

We conducted sensitivity analyses that assessed the associations between SES and hypertension (measured and medical) adjusted for potential confounders according to the new guidelines of ACC/AHA 2017, which were stratified by sex and urbanity (S2-4 Fig). These analyses produced estimates and trends that are very similar to those for primary analyses, which reinforce our findings that increasing SES is associated with an increased likelihood of having hypertension, which differed by sex and urbanity.

Our sensitivity analyses constructed nested logistic regression models for the associations between blood pressure outcomes (medical) and SES that progressively adjusted for age, sex, urbanity, marital status, exposure to second-hand smoke and BMI (Table 4, S7 Table). The estimates and trends reinforce our primary findings.

For our secondary outcome, we examined the associations between SES and BMI using two different approaches. Firstly, we conducted sex and urbanity stratified analyses for SES and both global and South Asia specific categories of BMI (S5-7 Fig; S5 Table). We also tested BMI as a continuous variable in association with SES due to the low prevalence of obesity (S6 Table). These results supported that the likelihood of being overweight/obese increased with increasing level of SES, which also differed by sex and urbanity.

## Discussion

Our study, including 13 436 people from a nationally representative survey, demonstrated that increasing levels of SES (education and wealth) were positively associated with an increased risk of having hypertension in Nepal, with strong evidence that men were more likely to be hypertensive than women. These associations also vary by urbanity. Our novel finding is that BMI partially mediated the associations between SES and hypertension in the context of LMICs. We found these results were comparable for both the JNC 7 and the ACC/ AHA 2017 guidelines.

Established evidence suggests that risk factors for CVD, including hypertension, are highly prevalent in low SES group in developed countries.^12,23^ In contrast to this evidence, our study shows that the prevalence of hypertension was greater among people with higher SES groups, which is consistent with recent studies conducted in LMICs,^24–26^ particularly in South Asia.^15,27^ Substantial differences between male and female were observed in the association between SES and hypertension, which is consistent with previous studies in developed countries.^15,28,29^

In line with these studies, our study observed that increasing levels of education and wealth quintiles have a positive association with higher likelihood of BMI both in men and women. Hence, we formally tested the mediating roles of BMI in the association between higher SES and hypertension and demonstrated that BMI attenuates the observed associations. In other words, BMI may help to explain broader SES differentials in hypertension, particularly by education and wealth quintiles. Evidence from higher income countries also supported that BMI mediates the association between education and the risk of cardiovascular diseases.^32^

Our observed results have several policy implications. The comprehensive understanding of the mechanisms of socioeconomic differentials in hypertension may help to take effective measures for the prevention of risk of CVD in resource-poor settings. Findings related to SES by sex differences in hypertension will also guide to take gender-sensitive policy measures in reducing CVD and its modifiable risk factors.

Moreover, the identification of BMI as a mediator of the higher SES and hypertension association, emphasize to this modifiable risk factor as a potential target for interventions to reduce CVD and related risk factors such as hypertension and elevated blood pressure in higher SES groups in LMICs. This study provides further evidence allied to the emergence of SES gradients in CVD and related risk factors. Although few recent studies found SES gradients in CVD risk in LMICs setting, this research contributes to previous work by bridging the fields of socioeconomic differentials in CVD risk and formally testing established theoretical models. The veracity of our findings is contingent on replication with longitudinal data and more comprehensive assessments of SES.

To the best of our knowledge, this is the first study found mediating roles of the modifiable risk factors of CVD in the SES and hypertension association using a nationally representative sample in a resource-poor setting. Our study also first time assessed the association between SES and hypertension according to standard hypertension JNC7 guideline and a new guideline recommended by the ACC/ AHA 2017. We observed sex and rural-urban differences in blood pressure outcomes by sex and urbanity stratified analysis. Furthermore, we have included medication adherence and multiple SES factors, which suggest an inclusive explanation of the association between higher SES and risk factors of CVD. For instance, recent studies also emphasized to investigate SES gradient along with sex and rural-urban differences in blood pressure outcomes in Nepal.^7, 8, 11, 33^

Along with these novel contributions and methodological strengths, some limitations may also be considered with the interpretation of the results. We were not able to assess the causality of the associations between SES and hypertension due to the cross-sectional nature of the data. Our measurement of SES omits an indicator of employment status which should be assessed in detail in further research. Blood pressure measurement error may occur due to the quality of medical staff training in various regions of Nepal even though an automatic device for BP measurement had been used. Finally, we were not able to assess the gender inequalities in access to medications use due to the insufficient information.

## Conclusions

In conclusion, higher SES was positively associated with the higher likelihood of having hypertension in Nepal according to both JNC7 and ACC/AHA 2017 guidelines. All of the observed trends were more pronounced in men than in women, and there was evidence of differences in these trends between residents in rural and urban areas. The association between higher SES and hypertension was mediated by BMI, which may help to explain broader socioeconomic differentials in CVD and related risk factors, particularly in terms of education and wealth. Our study suggests that the mediating factor of BMI should be tackled to diminish the risk of CVD in people with higher SES in LMICs.

## Acknowledgments

The authors thank to MEASURE DHS for granting access to the Nepal Demographic and Health Survey 2016 data. We also thank Prof. Parisa Tehranifar for her critical comments and suggestions on the initial version.

## Authors’ contributions

JR and RI developed the study concepts. JR, KKS, and RI analyzed the data. JR, ZA, KKS, and SB drafted the manuscript. All authors critically reviewed the manuscript. All authors have read and approved the final version of the paper.

## Ethical Approval and Consent to participate

The survey protocol for the primary data collection was approved by the ICF Institutional Review (IRB) and the Ministry of Health Ramshah Path, Kathmandu, Nepal. Informed consent was taken from each participant before the enrollment in the study. DHS program gave us formal approval to obtain the de-identified data from the DHS online archive after. Details: https://dhsprogram.com/data/Using-325 DataSets-for-Analysis.cfm

## Supporting information

### Supplementary Tables

**S1 Table. Association of blood pressure outcome with education levels stratified by sex in Nepal.** Odds ratios are adjusted for age, urbanity and marital status; * p<0.05, ** p<0.01, *** p<0.001.

**S2 Table. Association of blood pressure outcome with wealth quintiles stratified by sex in Nepal.** Odds ratios are adjusted for age, urbanity and marital status; * p<0.05, ** p<0.01, *** p<0.001.

**S3 Table. Association of blood pressure outcome with education levels stratified by urbanity in Nepal.** Odds ratios are adjusted for age, sex and marital status; * p<0.05, ** p<0.01, *** p<0.001.

**S4 Table. Association of blood pressure outcome with wealth quintiles stratified by urbanity in Nepal.** Odds ratios are adjusted for age, sex and marital status; * p<0.05, ** p<0.01, *** p<0.001.

**S5 Table. Association of Overweight/Obesity (South Asia specific definition) with education levels and wealth quintiles by urbanity in Nepal.** Model 1: Association of Overweight-SA with Education Level (Odds are adjusted for age, marital status and urbanity; * p<0.05, ** p<0.01, *** p<0.001). Model 2: Association of Overweight-SA with Wealth Quintiles (Odds are adjusted for age, marital status and urbanity; * p<0.05, ** p<0.01, *** p<0.001).

**S6 Table. Linear Regression comparing BMI with educational levels and wealth quintiles in Nepal.** Model 1: Association between BMI and Education Levels (Coefficients are adjusted for age, marital status, and urbanity; * p<0.05, ** p<0.01, *** p<0.001). Model 2: Association between BMI and Wealth Quintiles (Coefficients are adjusted for age, marital status and urbanity; * p<0.05, ** p<0.01, *** p<0.001).

**S7 Table. Mediation effect of BMI on Normal Blood Pressure Outcome (medical) by Education Levels and Wealth Quintiles.** aCoefficients adjusted for age, sex, marital status, urbanity, and second-hand smoking; bCoefficients further adjusted for body mass index (BMI); Exponentiated coefficients; 95% confidence intervals in brackets * p<0.05, ** p<0.01, *** p<0.001.

### Supplementary Figures

**S1 Fig.** Flow chart of analytic sample selection process

**S2 Fig. Association of hypertension (measured and medical) by ACC/AHA 2017 guideline with education levels by sex in Nepal.** Odds ratios are adjusted for age, urbanity and marital status, and stratified by sex. Measurement-only outcomes are defined based on cut-off points only; medical outcomes are defined based on cut-off points, diagnosis by a health professional or relevant medication use.

**S3 Fig. Association of hypertension (measured and medical) by ACC/AHA 2017 guideline with wealth quintiles by sex in Nepal.** Odds ratios are adjusted for age, urbanity and marital status, and stratified by sex. Measurement-only outcomes are defined based on cut-off points only; medical outcomes are defined based on cut-off points, diagnosis by a health professional or relevant medication use.

**S4 Fig. Association of hypertension (medical) by ACC/AHA 2017 guideline with education and wealth quintiles by urbanity in Nepal.** Odds ratios are adjusted for age, sex, and marital status, and stratified by urbanity. Medical outcomes are defined based on cut-off points, diagnosis by a health professional, or relevant medication use.

**S5 Fig. Association of overweight/obese with a) education levels b) wealth quintiles by the sex in Nepal.** Odds ratios are adjusted for age, urbanity and marital status, and stratified by sex. Overweight and obese are defined as BMI ≥ 23 kg/m2 (South Asia-specific definitions).

**S6 Fig. Association of overweight/ obese (global definition) with a) education levels b) wealth quintiles by the sex in Nepal.** Odds ratios are adjusted for age, urbanity and marital status and stratified by sex. Overweight and obese are defined as BMI ≥ 25 kg/m2 (Global definitions).

**S7 Fig. Association of overweight/ obese (global definition) with a) education levels b) wealth quintiles by the urbanity in Nepal.** Odds ratios are adjusted for age, sex, and marital status and stratified by the place of residence. Overweight and obese are defined as BMI ≥ 25 kg/m2 (Global definitions).

